# Establishment of age-specific intervals based on the hematological and immunological parameters of commercial pigs in the Republic of Korea

**DOI:** 10.1101/2024.12.18.626293

**Authors:** Won Gyeong Kim, Hansong Chae, Doyoung Song

## Abstract

Health monitoring based on hematological and immunological parameters is crucial for optimizing performance in the swine industry. This study was conducted to address the lack of comprehensive reports on these parameters in commercial pigs in the Republic of Korea (ROK). To fill this gap, blood samples from 764 healthy pigs across 32 farms in the ROK were analyzed, with the pigs categorized into specific age groups, including weaners, growers, finishers, gilts, and sows. Complete blood counts (CBC) and immunoglobulin G (IgG) concentrations were measured, revealing significant differences across all age groups (p<0.0001). These findings allowed us to establish age-specific Reference Intervals (RIs) at a 95% confidence level and Confidence Intervals (CIs) at a 90% confidence level for each group. This study provides robust guidelines for evaluating the health status of commercial pigs at different stages of growth and development in the ROK. The established RIs and CIs offer valuable insights for future research and practical applications, ensuring reliable health assessments. Routine monitoring of these parameters represents a critical tool for early detection and improved health management in commercial pig populations.

## Background

Pork is one of the most widely consumed meats, leading to extensive research to enhance pig productivity. Continuous disease control and immunity management are among the most effective strategies for maintaining pig health and achieving a sustainable swine industry. Such management practices help reduce economic losses and boost productivity.

The success of the swine industry is closely tied to the health and productivity of its livestock, making health monitoring through hematological and immunological analyses essential. Hematological analysis, in particular, is a powerful tool for detecting variations in blood parameters resulting from infections, inflammation, or other health concerns [1]. These parameters also serve as valuable markers for feed efficiency, growth rates, and reproductive performance, highlighting their importance in swine management [2-4].

To maximize the utility of hematological analyses for monitoring swine health, establishing Reference Intervals (RIs) and Confidence Intervals (CIs) from a healthy population is crucial. These intervals provide baseline values for assessing the health of individual pigs and herds, enabling early detection of indicators of disease or other health issues. Additionally, RIs and CIs offer consistent standards for distinguishing between normal variations and clinically significant changes in hematological values [5].

RIs and CIs are essential tools for informed decision-making, offering veterinarians and farm managers reliable indicators for determining whether intervention is required [1, 5]. By supporting consistent health monitoring, these intervals ensure that significant health deviations are addressed promptly, contributing to long-term herd management and improved productivity within the swine industry.

Furthermore, RIs and CIs can be tailored to account for variations in species, age, and environmental factors, ensuring precise interpretation of hematological data. This specificity not only enhances the reliability of collected data but also ensures accurate identification of health and disease states [1, 5]. Establishing these intervals improves swine health management, ultimately contributing to higher productivity.

Despite their significance, large-scale studies on hematological and immunological parameters in commercial pigs in the Republic of Korea (ROK) are limited. Previous research has primarily focused on weaners and miniature pigs for laboratory purposes, with little attention to commercial pig populations. Consequently, RIs from other countries are often applied to pigs in the ROK, which may lead to inaccuracies due to population-specific differences.

To address this gap, this study aimed to establish age-specific RIs and CIs for hematological and immunological parameters in a population of 764 clinically healthy commercial pigs. The pigs, categorized into nursery pigs, growers, finishers, gilts, and sows, were sampled from 32 farms across the ROK. These findings provide population-specific reference standards, enabling accurate health assessments and improved management practices for commercial pigs in the ROK.

## Methods and Material

### Sample collections

As part of the health monitoring program offered by Animal Industry Data Korea Corp. (AIDKR), a total of 764 blood samples were obtained from 32 farms in the ROK in 2021 and 2022.

To analyze the hematological and immunological parameters, veterinarians conducted clinical examination of the pigs and discussed the overall farm condition with farm workers before randomly selected clinically healthy animals. The blood samples from the selected pigs were collected by skilled veterinarians through jugular venipuncture. The samples were collected in appropriate tubes containing the anticoagulant ethylenediamine tetraacetic acid (EDTA) for a complete blood cell count (CBC), and serum separation tube (Becton, Dickinson and Company, NJ, USA) for serum collection. All collected samples were promptly transported to the AIDKR laboratory at a temperature of 4°C within 6 h.

### Hematological analyses

CBC was performed using the Prokan PE-6800VET analyzer (Shenzhen Prokan Electronics Inc., Shenzhen, China) following the manufacturer’s instructions. The CBC profile comprised a total of 18 parameters. The white blood cell (WBC) parameters included WBC count, lymphocyte count (LYM#) and its percentage (LYM%), and granulocyte count (GRAN#) and its percentage (GRAN%). The red blood cell (RBC) parameters included RBC count, hemoglobin concentration (HGB), hematocrit (HCT), mean corpuscular volume (MCV), mean corpuscular hemoglobin (MCH), mean corpuscular hemoglobin concentration (MCHC), red cell distribution width-standard deviation (RDW-SD), and red cell distribution width-coefficient of variation (RDW-CV). The platelet (PLT) parameters included PLT count, mean platelet volume (MPV), platelet distribution width (PDW), plateletcrit (PCT), and platelet-large cell ratio (P-LCR).

### Quantification of serum total immunoglobulin G (IgG)

For the immunological parameters, total IgG was quantified in serum using pig IgG enzyme-linked immunosorbent assay (ELISA) quantitation kits (Bethyl Laboratories Inc, Montgomery, USA). Following the manufacturer’s instructions, 100 µL of diluted affinity-purified antibody was coated on MaxiSorp plates (Thermo Fisher Scientific, Waltham, USA); antibody was diluted with 0.05 M carbonate-bicarbonate buffer at 1:100 ratio. The plates were then incubated for 1 h at 25°C and washed five times using 350 µL of washing buffer per well (0.05 M Tris + 0.14 M NaCl + 0.05% Tween 20). To block the plates, 200 µL of blocking buffer (0.05 M Tris + 0.14 M NaCl + 1.0% BSA; pH 8.0) was added to each well and incubated for 30 min at 25°C. After washing, 100 µL each of standard concentration and diluted serum were added to the plates and incubated for 1 h at 25°C. Horseradish peroxidase (HRP) detection antibody was diluted at 1:150,000 in a sample/conjugate diluent (0.05 M Tris + 0.14 M NaCl + 1.0% BSA + 0.05% Tween 20; pH 8.0). The plates were washed five times, and 100 µL of the HRP substrates were added and incubated for 1 h at 25°C. Following another round of washing, 100 µL of 3,3_′_,5,5_′_-tetramethylbenzidine (TMB) was added to each well and incubated in dark for 10 min at 25°C. Then, the colorimetric reaction was stopped by adding 100 µL of stop solution (0.18 M H_2_SO_4_), and the optical density (OD) was measured at 450 nm using an Infinite pro 200 microplate reader (Tecan, Männedorf, Switzerland). Total IgG concentration was calculated based on the 4-parameter logistic curve fit of the standard curve prepared using the OD values.

### Statistical analysis

All data were analyzed using GraphPad Prism version 9.0.0 (GraphPad Software, San Diego, California USA) and R version 4.2.2 (https://www.R-project.org/) with “reference Intervals” package version 1.2.0 (Finnegan D, https://CRAN.R-project.org/package=referenceIntervals). Horn’s test was performed for detecting outliers. Normality of the data for each dataset was evaluated using the Shapiro-Wilk test, with p<0.05 being considered statistically significant. Differences in each parameter among different age groups of pigs were determined using one-way ANOVA or Kruskal-Wallis test based on the data distribution, along with Dunn’s multiple comparisons test as a post-hoc; p<0.05 was considered statistically significant. Moreover, 95% RIs with 90% confidence intervals (CI) were determined based on the target sample size according to the RI guidelines for veterinary species [5].

## Results

### Sample collection information

We collected a total of 764 blood samples from 32 farms in the ROK between 2021 and 2022 (Supplementary Table 1). We divided pigs into five age categories: weaners, growers, finishers, gilts, and sows. Blood samples were collected from 95 weaners, 165 growers, 114 finishers, 177 gilts, and 213 sows.

### Values of hematological and immunological parameters in commercial pigs of various age groups in the ROK

To determine the normal range of hematological and immunological parameters for commercial pigs of various age groups, we analyzed 19 parameters in clinically healthy commercial pigs based on five age categories. All 19 parameters showed significant differences among different age groups, as shown in Table 1 (*p*<0.0001). Values of the leukocyte-related parameters—such as WBC count, LYM#, and GRAN#—were found to decrease with age. In contrast, LYM% increased with age and then decreased in sows, while GRAN% in sows significantly increased compared to other age groups. Values of erythrocyte-related parameters—RBC count and HCT—first increased and then decreased with age. Moreover, HGB, MCV, MCH, and MCHC also initially increased but then decreased with age, while RDW-SD and RDW-CV in sows were significantly different compared to other age groups. Values of thrombocyte-related parameters showed that PLT count decreased with age; MPV and PDW decreased and then increased with age. P-LCR increased and then decreased with age, and the PCT count in weaners was significantly higher than that of other age groups. Furthermore, despite fluctuations among the age groups, total IgG concentrations showed an overall increase with age.

**Table 1.** Comparison of hematological and immunological parameters between commercial healthy pigs of different age groups in ROK.

| Parameters* | Unit | Weaners<br>(N=95) | Growers<br>(N=165) | Finishers<br>(N=114) | Gilts<br>(N=177) | Sows<br>(N=213) |
| --- | --- | --- | --- | --- | --- | --- |
| WBC | 10<sup>3</sup>/uL | 20.1 ± 0.30 | 17.8 ± 0.17 | 17.0 ± 0.24 | 15.6 ± 0.23 | 12.6 ± 0.2 |
| LYM% | % | 25.2 ± 0.50 | 26.0 ± 0.28 | 28.5 ± 0.3 | 29.6 ± 0.4 | 23.4 ± 0.6 |
| GRAN% | % | 65.4 ± 0.24 | 64.9 ± 0.2 | 66.6 ± 0.2 | 65.7 ± 0.1 | 72.4 ± 0.2 |
| LYM# | 10<sup>3</sup>/uL | 5.0 ± 0.18 | 4.7 ± 0.1 | 4.8 ± 0.1 | 4.6 ± 0.1 | 3.0 ± 0.1 |
| GRAN# | 10<sup>3</sup>/uL | 13.1 ± 0.28 | 11.7 ± 0.2 | 11.1 ± 0.1 | 10.1 ± 0.2 | 9.0 ± 0.2 |
| RBC | 10<sup>6</sup>/uL | 5.8 ± 0.01 | 6.3 ± 0.01 | 6.4 ± 0.01 | 6.5 ± 0.02 | 5.6 ± 0.02 |
| HGB | g/dL | 9.7 ± 0.03 | 10.3 ± 0.02 | 11.3 ± 0.04 | 11.8 ± 0.0 | 11.2 ± 0.04 |
| HCT | % | 34.7 ± 0.10 | 37.4 ± 0.05 | 38.1 ± 0.1 | 38.8 ± 0.14 | 36.7 ± 0.11 |
| MCV | fL | 59.7 ± 0.05 | 59.6 ± 0.03 | 59.9 ± 0.05 | 60.8 ± 0.03 | 65.3 ± 0.06 |
| MCH | Pg | 16.5 ± 0.02 | 16.5 ± 0.01 | 17.6 ± 0.02 | 18.2 ± 0.01 | 19.7 ± 0.02 |
| MCHC | g/dL | 27.7 ± 0.01 | 27.5 ± 0.02 | 29.6 ± 0.02 | 30.1 ± 0.01 | 30.1 ± 0.02 |
| RDW-SD | fL | 35.5 ± 0.15 | 34.6 ± 0.12 | 35.6 ± 0.12 | 35.9 ± 0.11 | 37.0 ± 0.04 |
| RDW-CV | % | 18.4 ± 0.04 | 17.95 ± 0.03 | 18.53 ± 0.03 | 18.49 ± 0.02 | 17.81 ± 0.01 |
| PLT | 10<sup>3</sup>/uL | 398.0 ± 38.86 | 264.06 ± 18.58 | 247.43 ± 21.46 | 234.09 ± 13.86 | 175.03 ± 13.63 |
| MPV | fL | 9.5 ± 0.07 | 8.86 ± 0.02 | 8.94 ± 0.02 | 8.90 ± 0.02 | 9.7 ± 0.04 |
| PDW | % | 10.5 ± 0.13 | 9.70 ± 0.13 | 9.34 ± 0.05 | 9.38 ± 0.05 | 11.00 ± 0.11 |
| PCT | % | 0.4 ± 0.05 | 0.25 ± 0.02 | 0.25 ± 0.06 | 0.20 ± 0.02 | 0.16 ± 0.02 |
| P-LCR | % | 11.9 ± 0.59 | 12.59 ± 0.43 | 14.54 ± 0.64 | 13.29 ± 0.42 | 10.35 ± 0.36 |
| IgG | Mg/dL | 1041.4 ± 75.52 | 1328.60 ± 33.11 | 1186.11 ± 29.64 | 1262.38 ± 23.30 | 1487.64 ± 33.99 |

<sup>\*</sup> $p < 0.0001$ for all parameters.

### RIs and CIs for hematological and immunological parameters in commercial pigs of various ages groups in the ROK

We established 95% RIs and 90% CIs for the lower and upper reference limits of 19 hematological and immunological parameters in commercial pigs according to their age groups. The established RIs specific to each age group of commercial pigs are presented in Table 2.

**Table 2.** Reference intervals for hematological and immunological parameters of healthy pigs in ROK.

| Parameters | Weaners |  | Growers |  | Finishers |  | Gilts |  | Sows |  |
| --- | --- | --- | --- | --- | --- | --- | --- | --- | --- | --- |
| (Unit) | 95% RI | 90% CI | 95% RI | 90% CI | 95% RI | 90% CI | 95% RI | 90% CI | 95% RI | 90% CI |
| WBC<br>(10<sup>3</sup>/uL) | 11.6-30.9 | 10.6-32.7 | 10.5-24.8 | 9.3-26.3 | 10.0-25.9 | 9.1-27.3 | 9.1-24.3 | 8.4-26.0 | 7.0-19.5 | 6.4-21.6 |
| LYM<br>(%) | 13.2-40.9 | 11.8-43.6 | 16.8-39.9 | 16.0-46.2 | 18.7-40.2 | 17.5-42.0 | 16.8-42.9 | 13.7-44.6 | 10.3-39.8 | 8.3-45.8 |
| GRAN<br>(%) | 50.8-81.8 | 48.6-85 | 48.4-75.8 | 47.2-80.5 | 54.8-80.8 | 52.8-82.5 | 53.2-76.3 | 52.8-77.0 | 59.3-87.6 | 55.5-90.2 |
| LYM#<br>(10<sup>3</sup>/uL) | 2.0-9.3 | 1.7-10 | 2.5-8.1 | 2.0-8.9 | 2.4-7.9 | 2.2-8.4 | 2.4-8.0 | 1.8-8.9 | 1.3-7.0 | 0.9-7.8 |
| GRAN#<br>(10<sup>3</sup>/uL) | 6.7-2.6 | 5.9-23.1 | 6.1-17.6 | 5.1-20.1 | 6.9-16.2 | 6.4-17.0 | 5.5-15.7 | 5.1-17.8 | 4.9-14.5 | 4.5-16.0 |
| RBC<br>(10<sup>6</sup>/uL) | 4.9-6.9 | 4.7-7 | 5.4-6.9 | 5.2-7.2 | 5.5-7.4 | 5.3-7.5 | 5.1-7.8 | 5.0-8.1 | 4.1-7.1 | 3.9-7.2 |
| HGB<br>(g/dL) | 7.7-11.9 | 7.7-12.9 | 8.4-12.6 | 8.1-12.8 | 9.0-14.0 | 8.6-14.3 | 8.5-14.6 | 8.0-15.5 | 8.3-13.9 | 7.7-14.3 |
| HCT<br>(%) | 27.8-42.4 | 26.8-43.6 | 31.3-43.8 | 30.6-44.4 | 31.0-45.9 | 30.1-47.0 | 28.3-48.0 | 27.7-49.5 | 28.6-45.0 | 26.7-46.0 |
| MCV<br>(fL) | 52.9-66.9 | 52.0-68.0 | 53.6-65.9 | 52.8-67.3 | 53.3-66.8 | 52.5-67.7 | 56.1-67.2 | 54.8-68.2 | 59.2-73.0 | 57.6-74.5 |
| MCH<br>(Pg) | 14.5-18.5 | 14.3-18.8 | 14.7-18.5 | 14.1-18.7 | 15.7-19.7 | 15.4-20.0 | 16.4-20.1 | 15.9-21.3 | 17.4-22.1 | 17.1-23.1 |
| MCHC<br>(g/dL) | 25.4-30.2 | 25.0-30.5 | 23.9-30.5 | 23.5-31.1 | 26.6-32.7 | 26.2-33.2 | 27.8-32.3 | 27.6-33.1 | 27.5-33.1 | 27.2-33.6 |
| RDW-SD<br>(fL) | 27.0-45.1 | 25.9-46.6 | 27.9-42.4 | 24.1-44.6 | 27.9-44.7 | 27.1-46.1 | 27.9-42.7 | 27.9-44.6 | 32.0-42.7 | 31.6-42.7 |
| RDW-CV<br>(%) | 15.2-21.8 | 14.8-22.3 | 15.2-21.0 | 15.1-21.6 | 15.8-21.5 | 15.4-21.9 | 15.7-21.3 | 15.3-21.4 | 16.0-19.9 | 15.6-20.4 |
| PLT<br>(10<sup>3</sup>/uL) | 52.7-1037.0 | 32.3-1157.7 | 71.1-693.9 | 58.0-813 | 44.2-615.3 | 29.0-679.3 | 73.3-563.7 | 60.0-679.0 | 39.6-427.8 | 34.0-473.0 |
| MPV<br>(fL) | 6.2-12.6 | 5.7-13.4 | 7.7-11.5 | 7.6-12.5 | 7.2-10.8 | 7.0-11.1 | 7.7-10.7 | 7.1-12.5 | 7.5-12.1 | 7.4-13.3 |
| PDW<br>(%) | 6.2-15.7 | 5.8-16.8 | 7.7-15.4 | 7.4-15.4 | 6.5-12.1 | 6.2-12.6 | 7.7-12.7 | 7.4-13.3 | 7.7-15.4 | 7.7-15.4 |
| PCT<br>(%) | 0.02-1.16 | 0.004-1.32 | 0.1-0.9 | 0.1-1.0 | 0.03-0.60 | 0.02-0.67 | 0.04-0.65 | 0.04-0.79 | 0.03-0.45 | 0.02-0.5 |
| P-LCR<br>(%) | 3.2-23.7 | 2.5-26.5 | 6.2-29.3 | 5.4-31.8 | 4.6-28.7 | 3.9-31.4 | 7.1-27.3 | 6.0-33.2 | 4.8-22.7 | 4.3-25.3 |
| IgG<br>(Mg/dL) | 232.3-2431.2 | 162.2-2684.6 | 687.0-2410.3 | 588.3-2758.9 | 562.9-2051.2 | 499.0-2166.5 | 596.5-2055.6 | 559.4-2121.0 | 788.9-3034.0 | 725.6-3633.1 |

## Discussion

In the field of swine health management, RIs and CIs for hematological and immunological parameters help ensure the accuracy of experimental data by identifying potential errors and are useful guidelines for veterinarians to make accurate diagnoses, make treatment, and monitor the health of farm-raised pigs.

RIs serve as indicators for determining whether data falls within the normal range [1, 4, 5]. For example, if a WBC count is located within RI, it can be considered that the WBC count of the pig is within the expected range for its age. Consequently, the RIs can provide a reference range for assessing whether the WBC count is normal. CIs, on the other hand, indicate that the possibility that the data corresponds to the population. If a WBC count located within the CI, it suggests that the WBC count of the pig being like the average value in the same age group. This provides a range of confidence regarding the accuracy of the data. However, for commercial pigs in the ROK, reliable and sufficient data on these parameters have been scarce due to the cost- and time-intensive nature of comprehensive assessments. Therefore, this study aimed to assess the values of these parameters and establish RIs and CIs specific to commercial pigs in the ROK.

We conducted experiments using blood from clinically healthy pigs and established RIs and CIs for hematology and immunology based on the experimental data, according to different growth stages (Table 2). In this study, significant differences were observed in all hematological and immunological parameters when comparing different age groups of commercial pigs. When compared with studies from other research, it was consistent that there were differences in RIs and CIs across each age group.

These findings highlight the need to establish RIs specific to each age group in order to precisely interpret the health status of commercial pigs in the ROK. Similar observations have been reported in previous studies. For instance, a study in China found significant differences in 19 out of 25 hematological parameters between commercial weaners and sows, leading to the establishment of separate RIs for each group [6]. Similarly, in Slovenia, significant differences were observed in 11 out of 14 hematological parameters between commercial growers and finishers, as well as in 14 out of 20 hematological parameters among growers, finishers, and sows, necessitating the establishment of distinct RIs for each category [7, 8].

In addition, within the same age group, the RIs and CIs values differed between studies. For instance, in this study, the RI for WBC count in weaners was 11.6-30.9 × 10^3^/µL, with CIs of 10.6-32.7 × 10^3^/µL. In comparison, the values reported in China were 9.34-23.84 × 10^3^/µL (CIs: 8.26-24.92 × 10^3^/µL), and in Canada, 6.0-21.7 × 10^3^/µL (CIs: 5.8-22.8 × 10^3^/µL) [1, 6]. We thought the following reasons for the differences in RIs and CIs studies for hematological parameters across various research studies: these differences are likely influenced not only by variations in pig species but also by farming and environmental conditions.

However, hematological values can significantly vary due to several factors such as age, gender, and physiological characteristics [6]. Especially, the quality of the blood can affect the results, such as storage temperature, coagulation and hemolysis. Therefore, if data from a clinically healthy pig appears abnormal results or is suspected to be an outlier within the same herd, these factors should be considered as blood quality issue. Thus, these factors can lead to the misinterpretation of whether abnormal results are due to illness or poor blood quality.

Given the similarities with previous studies and the comprehensive assessment of hematological and immunological parameters across all age groups of commercial pigs, the significant differences observed in all evaluated parameters in this study were considered reasonable. Furthermore, these observations emphasized the importance of establishing age-specific RIs and CIs for the effective management of commercial pigs in the ROK. Additionally, since there have been no previous studies on RIs for finishers and gilts, our results could be valuable for veterinarians and other researchers.

## Conclusions

In conclusion, we observed that hematology parameters exhibit variability during the growth of commercial pigs and therefore established age-specific RIs and CIs of hematological and immunological parameters for commercial pigs in the ROK.

Our findings were similar in verifying that RI and CI values differ at each growth stage, and we confirmed that even at the same growth stage, RI and CI values vary between different countries. This suggests the importance of age groups in influencing hematological parameters and indicates that differences in hematological parameters may be influenced by factors such as pig breeds and rearing environments. This difference in RIs and CIs values growth stages suggests the necessity of conducting routinely tests at each stage to effectively manage health of pigs.

Additionally, since the RIs and CIs presents in this study are based on clinically healthy pigs and appropriate experimental conditions, they can serve as valuable references for evaluating hematological and immunological results in commercial pigs in the ROK.

Given the substantial number of commercial pigs and domestic farms in the ROK, further studies with larger sample sizes are necessary to establish more reliable RIs and CIs for hematological and immunological parameters. This will enhance our understanding of the normal range of these parameters in commercial pigs, facilitating effective health monitoring and diagnosis of potential abnormalities or diseases.

## List of abbreviations

AIDKR: Animal Industry Data ROK Corp.
CBC: complete blood cell count
CI: confidence interval
EDTA: ethylenediamine tetraacetic acid
ELISA: enzyme-linked immunosorbent assay
GRAN: granulocyte
HCT: hematocrit
HGB: hemoglobin concentration
HRP: Horseradish peroxidase
IgG: immunoglobulin G
LYM: lymphocyte
MCH: mean corpuscular hemoglobin
MCHC: mean corpuscular hemoglobin concentration
MCV: mean corpuscular volume
MPV: mean platelet volume
OD: optical density
PCT: plateletcrit
PDW: platelet distribution width
P-LCR: platelet-large cell ratio
PLT: platelet
RBC: red blood cell
RDW-CV: red cell distribution width-coefficient of variation
RDW-SD: red cell distribution width-standard deviation
RI: reference interval
CI: confidence interval
WBC: white blood cell

## Declarations

### Ethics approval and consent to participate

This article did not require IRB/IACUC approval because it aimed to report the results retrospectively from the health monitoring service provided by Animal Industry Data Korea Corp.

### Consent for publication

Not applicable.

### Availability of data and materials

The datasets used and/or analyzed during the current study are available from the corresponding author on reasonable request.

### Authors’ contributions

DS conceptualized the study. DS and WK designed the methodology. WK and HC performed data curation. WK conducted formal analysis of data. HC validated the study. WK conducted investigations. DS, WK and HC wrote the original draft of this manuscript, and HC reviewed and edited the manuscript. All authors read and approved of the final manuscript.

## Acknowledgments

*-*

