## Supplementary table 1 for "Establishment of age-specific intervals based on the hematological and immunological parameters of commercial pigs in the Republic of Korea"

**Supplementary Table 1.** The number of samples collected per farm and age group for analyzing hematological and immunological parameters

| Farm | Weaners | Growers | Finishers | Gilts | Sows | Total |
| --- | --- | --- | --- | --- | --- | --- |
| 1 |  |  |  |  | 7 | 7 |
| 2 |  |  | 18 | 66 | 21 | 105 |
| 3 |  |  |  | 4 | 5 | 9 |
| 4 |  |  |  | 3 |  | 3 |
| 5 |  |  |  | 9 | 4 | 13 |
| 6 |  |  |  |  | 6 | 6 |
| 7 |  | 4 |  |  |  | 4 |
| 8 | 7 | 52 |  |  |  | 50 |
| 9 |  | 4 |  |  | 17 | 21 |
| 10 | 28 | 21 | 6 | 2 | 5 | 62 |
| 11 |  |  | 23 | 38 | 13 | 74 |
| 12 |  |  | 5 |  | 2 | 7 |
| 13 | 7 | 19 |  |  |  | 26 |
| 14 |  | 4 | 5 |  |  | 9 |
| 15 |  |  |  | 2 | 2 | 4 |
| 16 |  | 13 |  |  |  | 22 |
| 17 |  | 12 |  |  |  | 12 |
| 18 | 11 | 3 |  |  |  | 14 |
| 19 | 17 |  |  |  |  | 17 |
| 20 | 8 |  |  |  | 6 | 14 |
| 21 |  |  | 16 |  |  | 16 |
| 22 |  |  |  | 13 | 4 | 17 |
| 23 |  |  | 6 | 6 | 11 | 23 |
| 24 |  |  | 7 | 8 | 15 | 30 |
| 25 |  |  |  |  | 4 | 4 |
| 26 |  | 5 |  |  |  | 5 |
| 27 |  |  | 2 |  | 3 | 5 |
| 28 |  |  | 7 |  | 7 | 14 |
| 29 | 7 | 17 |  |  |  | 24 |
| 30 | 7 | 1 | 7 | 14 | 60 | 89 |
| 31 |  |  |  | 6 | 4 | 10 |
| 32 | 3 | 10 | 12 | 6 | 17 | 48 |
| Total | 95 | 165 | 114 | 177 | 213 | 764 |
